# Structure of far-red allophycocyanin: stripped down and tuned up for low energy photosynthesis

**DOI:** 10.1101/2025.02.25.640088

**Authors:** Giovanni Consoli, Ho Fong Leong, Geoffry A. Davis, Tom Richardson, Aiysha McInnes, James W. Murray, Andrea Fantuzzi, A. William Rutherford

## Abstract

A diverse subset of cyanobacteria is capable of transiently modifying their photosynthetic machinery in a process known as far-red light photoacclimation to drive photosynthesis with less energetic photons (700 nm - 800 nm). To achieve this all the main light-driven components of the photosynthetic apparatus, including their allophycocyanin antenna, are replaced with red-shifted paralogues. Recent studies based on the structure of an incomplete complex provided some insights into the tuning of the far-red phycobiliproteins. Here, we solved the structure of the intact bicylindrical allophycocyanin complex from the cyanobacterium *Chroococcidiopsis thermalis* PCC 7203 at a resolution of 2.61 Å determined by Cryo-electron microscopy single particle analysis. A comparison between far-red and white light allophycocyanin cores provides insight on the evolutionary adaptations needed to optimize excitation energy transfer in the far-red light adapted photosynthetic apparatus. The reduction in antenna size in far-red photosynthesis, suggests a need to optimize membrane packing to increase the number of photosystems, while the wider spread in the absorption range of the bilins suggests faster and more efficient excitation energy transfer to far-red Photosystem II by limiting backflow of excitation from the reaction centres to the far-red bilin pigments.

## Introduction

In some ecological niches, such as those found in cyanobacterial mats (Ohkubo & Miyashita, 2017), porous rocks (Trampe & Kühl, 2016), limestone caves (Behrendt et al., 2020) and other shaded environments (Antonaru et al., 2020), the ratio between visible light (400 nm – 700 nm) and far-red light (700 nm – 800 nm) decreases drastically. In these conditions, a small but phylogenetically diverse group of cyanobacteria is capable of far-red light photoacclimation (FaRLiP) (Gan et al., 2014), extending the “red limit” of photosynthesis using of photons up to 800 nm in wavelength (Nürnberg et al., 2018).

During this process, a gene cluster of ∼21 genes (Zhao et al., 2015), coding for specialized far-red-light-adapted paralogues of all the main components of the photosynthetic apparatus, replace their white light counterparts. The far-red paralogous subunits of Photosystem II (PSII) and Photosystem I (PSI) incorporate a small number of longer wavelength chlorophylls, Chlorophyll *d* (Chl *d*) and Chlorophyll *f* (Chl *f*) (∼8% to 10%) (Nürnberg et al., 2018; Chen et al., 2010) in specific positions, further redshifting their absorption spectra. On the other hand, the far-red paralogues of the phycobilisomes, the major light-harvesting antenna proteins in cyanobacteria, retain the same type of pigment, phycocyanobilin, shifting their absorption by modifying their protein environment (Soulier & Bryant, 2021; Gisriel et al., 2024; Ho et al., 2017).

Phycobilisomes are a structurally and functionally diverse family of light-harvesting complexes which bind open-chain tetrapyrroles, known as bilins, which absorb visible light and funnel excitation energy to the photosystems. The extreme variability in size, arrangement and absorbance profile of phycobilisomes reflects the evolutionary adaptation of these antenna complexes to a number of ecological niches (Bryant & Gisriel, 2024).

Given the small number of long wavelength chlorophylls present in far-red-adapted photosystems (Nürnberg et al., 2018), a red-shifted antenna, which is capable of efficiently funneling the excitation energy of far-red photons into the far-red photosystems, is necessary to increase the antenna size for wavelengths above 700 nm. The FaRLiP gene cluster contains five phycobiliprotein paralogues (ApcB2, ApcD5, ApcD3, ApcD2 and ApcE2), that assemble with other subunits in common with white light phycobilisomes (ApcF and ApcC) into a bicylindrical complex (Ho et al., 2017), which is significantly smaller than all the other classes of phycobilisomes (Bryant & Gisriel, 2024).

Similar to white-light phycobilisome (WL-PBS) cores, in far-red allophycocyanin (FR-APC), α- (ApcD2, ApcD3, ApcD5) and β- (ApcB2) heterodimers (αβ) assemble into toroid hexameric (αβ)_3_ complexes (Bryant, Glazer & Eiserling, 1976; Brejc et al., 1995; Gisriel et al., 2024) that are stacked with the help of ApcC onto a specialized core-membrane linker protein, ApcE2, that also anchors them to the stromal side of PSII (Zlenko et al., 2019). Compared to the far-red light-adapted subunits, the chromophores in the β-subunits (ApcB2) appear to be structurally more similar to its WL version (Gisriel et al., 2024) and therefore have similar absorption properties. On the other hand, in the α-subunits the pyrrole ring A of each bilin pigment is more coplanar with its other rings, leading to a red-shifted absorption compared to WL α-subunits, a feature that has also been observed in heterologously expressed biliproteins (Ho et al., 2017).

In the bicylindrical FR-APC, the bilins contained in ApcE2 and ApcD3 have been suggested to act as terminal emitters (Gisriel et al., 2024). In contrast to all other phycobiliproteins, these two subunits lack the cysteine that normally covalently binds the bilins. The absence of the covalent bond results in an extension of the π-system and a redshift of the absorption (Gan et al., 2014; Gan & Bryant, 2015; Miao et al., 2016; Xu et al., 2017). However, 27% of the ApcD3 sequences (to date) retain the bilin-binding Cys^78^ residue (Gan & Bryant, 2015; Gisriel et al., 2024), suggesting a degree of species specificity in the excitation energy transfer pathways to the photosystems via ApcD3.Click or tap here to enter text.

A previously determined CryoEM map of a disconnected portion of the FR-APC core complex from *Synechococcus* PCC 7335 contained only three of the four expected phycobiliprotein trimers (Gisriel et al., 2024). In the present work, we attempted the purification of a FR-APC + FR-PSII complex and obtained a structure of an intact bicylindrical FR-APC at a resolution of 2.61 Å, with a low-resolution ESP density localized where PSII is expected to bind. We describe the structure of the intact FR-APC complex, provide an analysis of the planarity of bilin pigments in relationship to their absorption, and suggest specific evolutionary traits needed to optimize far-red light excitation energy transfer.

## Results

### CryoEM data processing and map description

The CryoEM map of FR-APC was refined with C2 symmetry at a GS-FSC resolution of 2.61 Å (fig. S1, table S1). The map presents density to fit two slightly tilted, antiparallel far-red light adapted allophycocyanin cores (Fig. 1A). Each of the cylindrical allophycocyanin cores contains four (αβ)_3_ trimers, arranged in two face-to-face [(αβ)_3_]_2_ hexamers.

**Fig. 1.**
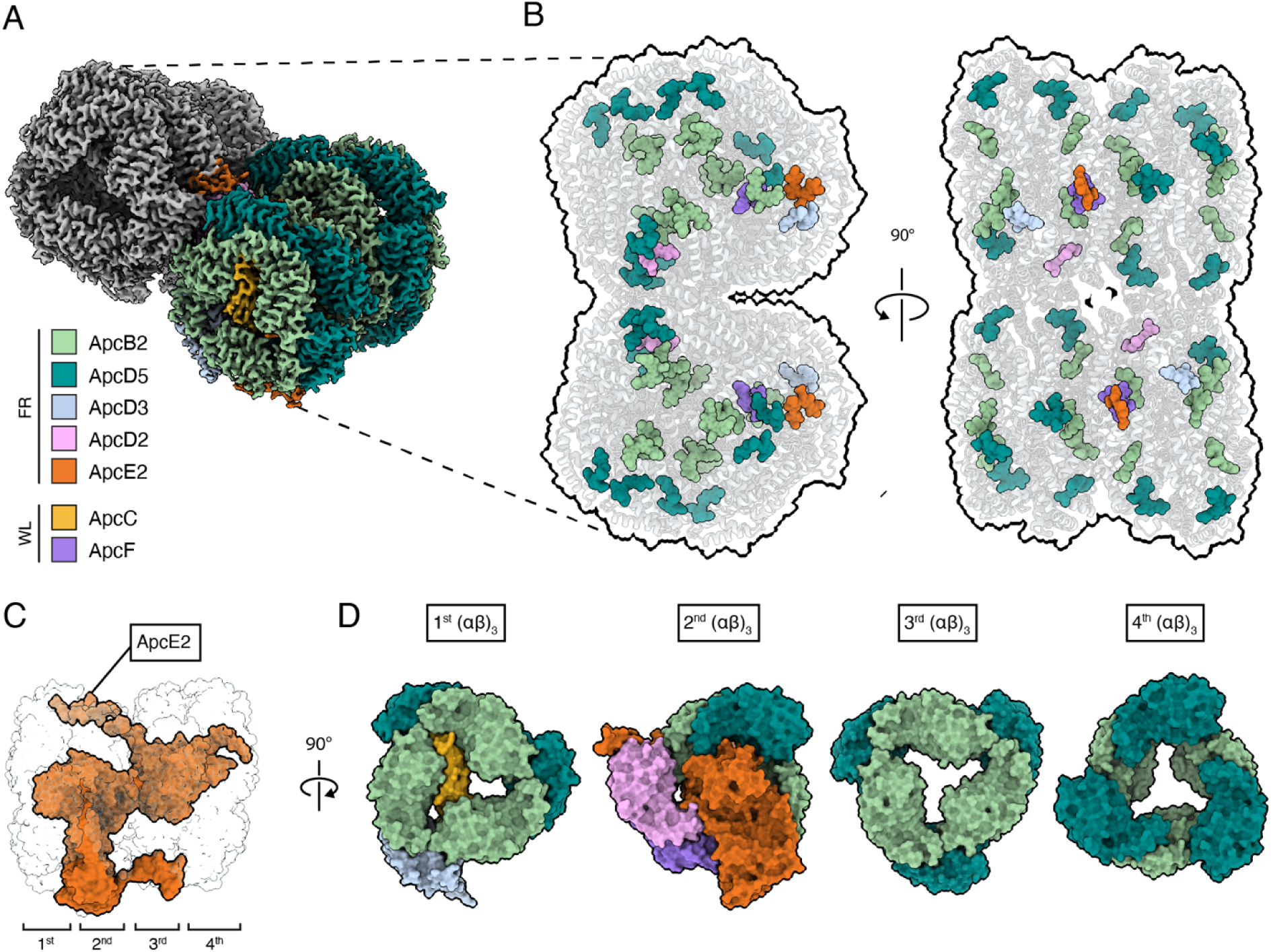
Structure of the bicylindrical FR-APC and positions of the bilin pigments. A) View of the dimeric FR-APC map from a tilted perspective facing the 1^st^ (αβ)_3_ from the cytoplasmic side. The map presents clear ESP for all the known FR-APC subunits, color key at the bottom left of the panel. B) Location of the phycocyanobilin pigments in the bicylindrical FR-APC complex as viewed from the membrane plane and from the cytosolic side of the membrane. The pigments are also colored according to subunit coordination in panel A. C) Side view of a single FR-APC cylinder with ApcE2 colored in orange and every other subunit transparent. D) Separate representation of the four (αβ)_3_ trimers, also colored according to the color key in panel A. It is important to note that the ApcE2 subunit is only represented in the second trimer.

The 1^st^ and 2^nd^ (αβ)_3_ trimers, which are proximal to the thylakoid membrane plane, contain the far-red light specific subunits ApcB2, ApcD5, ApcD3, ApcE2, ApcD2 and the white light shared subunits ApcC and ApcF (Fig. 1D). As suggested previously (Gisriel et al., 2024), the second hexamer (formed by the 3^rd^ and 4^th^ (αβ)_3_ trimers) is held in position by the second REP domain (REPeat domain) of ApcE2 and contains 6 ApcD5 and 6 ApcB2 subunits (Fig. 1D), but contrary to expectations (Gisriel et al., 2024), the distal ApcC subunit is not present. Due to its location, between 3^rd^ and 4^th^ (αβ)_3_ trimers, the absence of the distal ApcC is unlikely to be due its loss during isolation, since it would be sterically prevented from detaching by the presence of the fourth (αβ)_3_ trimer.

Even though some of the subunits (i.e. ApcB2 and ApcD5) are present in multiple copies in the structure, each bilin binding pocket is partially formed by the central subunits ApcE2 or ApcC, independently fine-tuning their chemical environment and consequently their absorption spectra. This can tune specific sites and model the site energy landscape of the pigments so that excitation can be efficiently funneled to the terminal emitter.

In addition, the map presents ESP density extending from the PB loop of ApcE2 that can be attributed to the presence of connected photosystems. Unfortunately, the limited number of particles available and heterogeneity prevent further local refinement of the photosystem portion of the map (fig. S1B).

### Absorption and low temperature fluorescence spectroscopy

The absorption spectrum of the FR-APC complex presents the same features previously reported in literature, with two main absorption peaks at 650 nm and 710 nm, corresponding to the β-subunits and α-subunits respectively (Gan et al., 2014; Ho et al., 2017). Moreover, the absorption spectrum presents, although with lower intensity, a peak around 440 nm, that indicates the presence of chlorophylls from photosynthetic complexes in the sample (Fig. 2A). Further analysis with low temperature fluorescence emission spectroscopy indicates that when chlorophylls are excited with blue light, there is a clear emission peak at 750 nm, with a shoulder at 730 nm, which can be interpreted as the fluorescence emission of FR-PSII, with some excitation being transferred back to the terminal emitter of the FR-APC. When exciting the sample with 550 - 600 nm light, where the only species absorbing are the bilin pigments, the fluorescence emission arises mainly from the bicylindrical FR-APC at 730 nm, with a smaller emission shoulder at 750 nm (Fig. 2B). This indicates that the sample is constituted mostly of disconnected FR-APC, with only a small proportion of the complexes being connected to FR-PSII.

**Fig. 2.**
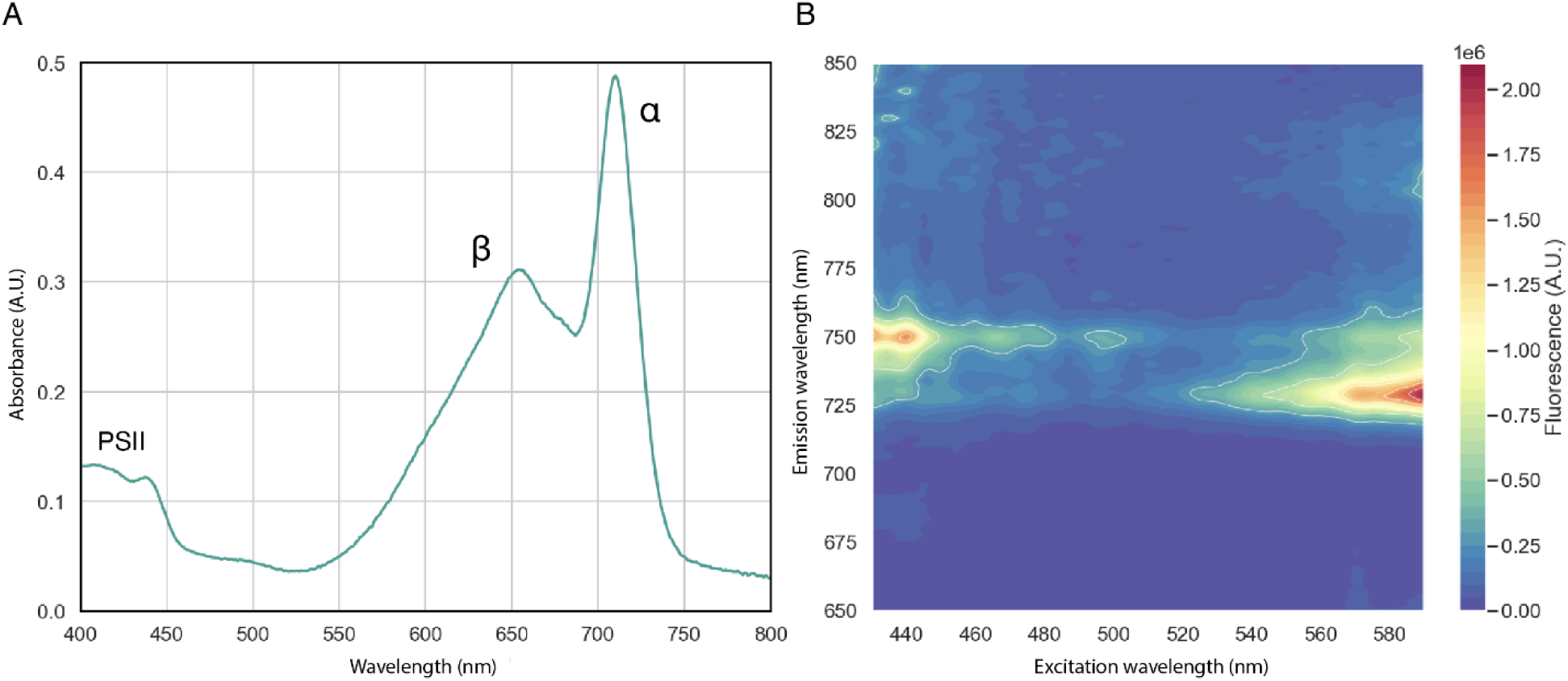
Absorption and fluorescence spectra of the bicylindrical FR-APC complex. A) Room temperature absorption spectra of the bicylindrical FR-APC, Absorbance peaks corresponding to PSII chlorophylls, β-subunits, and α-subunits are indicated. B) 2D low-temperature fluorescence (77 K) excitation *vs.* emission spectra of the bicylindrical FR-APC. The intensity of fluorescence is represented by the color gradient.

### Coordination of bilin sites

The planarity and chemical environment of the bilin pigments strongly influences their absorption spectra (Staheli et al., 2021; Soulier & Bryant, 2023). For this reason, to gain insight into the structural origin of the red-shifted absorbance of some of the subunits of the FR-APC complex (Fig. 2A), the angles between ring A and ring B of each bilin molecule and those between ring C and ring D were measured (Fig. 3). The angles between rings B and C do not vary significantly and are therefore not included in the analysis.

**Fig. 3.**
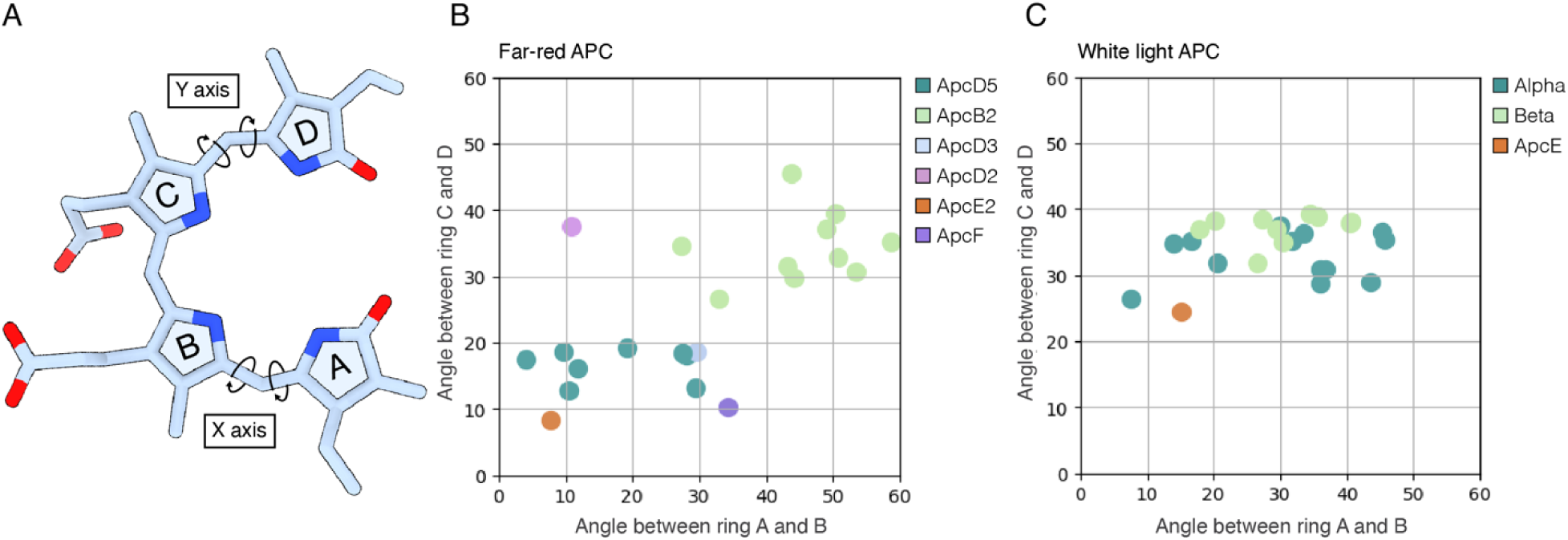
Analysis of the planarity of the bilin pigments in FR-APC and WL-APC. A) Representation of an unbound planar phycocyanobilin pigment with the measured torsion angles indicated. B) Variation of bilin planarity in FR-APC. The x-axis reports the degrees off plane of ring A with respect to ring B, while the y-axis reports the degrees off plane of ring D with respect to ring C. C) Variation of the planarity of the bilin pigments in WL-APC from *Synechocystis sp.* PCC 6803 (7SC7). The x-axis reports the degrees off plane of ring A with respect to ring B, while the y-axis reports the degrees off plane of ring D with respect to ring C.

The bilins contained in the α and β subunits of WL-APC (Domínguez-Martín et al., 2022) present very similar conformational landscapes, with the D ring being generally between 20° to 40° off plane and the A ring presenting a larger spread of 10° to 50° degrees off plane (Fig. 3B). The bilins contained in FR-APC present a more diverse arrangement, with the β-subunits (ApcB2) maintaining similar torsion angles compared with WL in the D ring, while the α subunits (i.e. ApcD2, ApcD3 and ApcD5) are far more planar than their WL counterparts (Fig. 3C). The effect of this change can be seen in the absorption spectra of the FR-APC complex (Fig. 2A), where the two main absorption peaks can be attributed to the less planar β-subunits (∼655 nm peak) and to the more planar α-subunits (∼710 nm peak) (Gisriel et al., 2024; Soulier, Laremore & Bryant, 2020).

Previous structures reported the loss of cysteine bonding in the subunits ApcE2 (Fig. 4) and ApcD3 (Fig. 5D), and therefore it was suggested that these two subunits contain the most red-shifted pigments which transfer excitation to FR-PSII (Gisriel et al., 2024). However, given that the loss of cysteine linkage in ApcD3 is only found in a few species (Ho et al., 2017) and that FR-PSII subunits are well conserved amongst each other, it is likely that the excitation injection point of FR-PSII from FR-APC is also conserved. This therefore suggests that ApcE2 is the main terminal emitter, also consistent with the bilin in ApcE2 being the most planar (Fig. 3B). ApcE2 has been suggested to have a key role in chlorophyll *d* synthesis (Bryant et al., 2020). Nonetheless, in the structure there are no other significant changes that would indicate an additional site for the synthesis of chlorophyll *d* by the FR-APC directly.

**Fig. 4.**
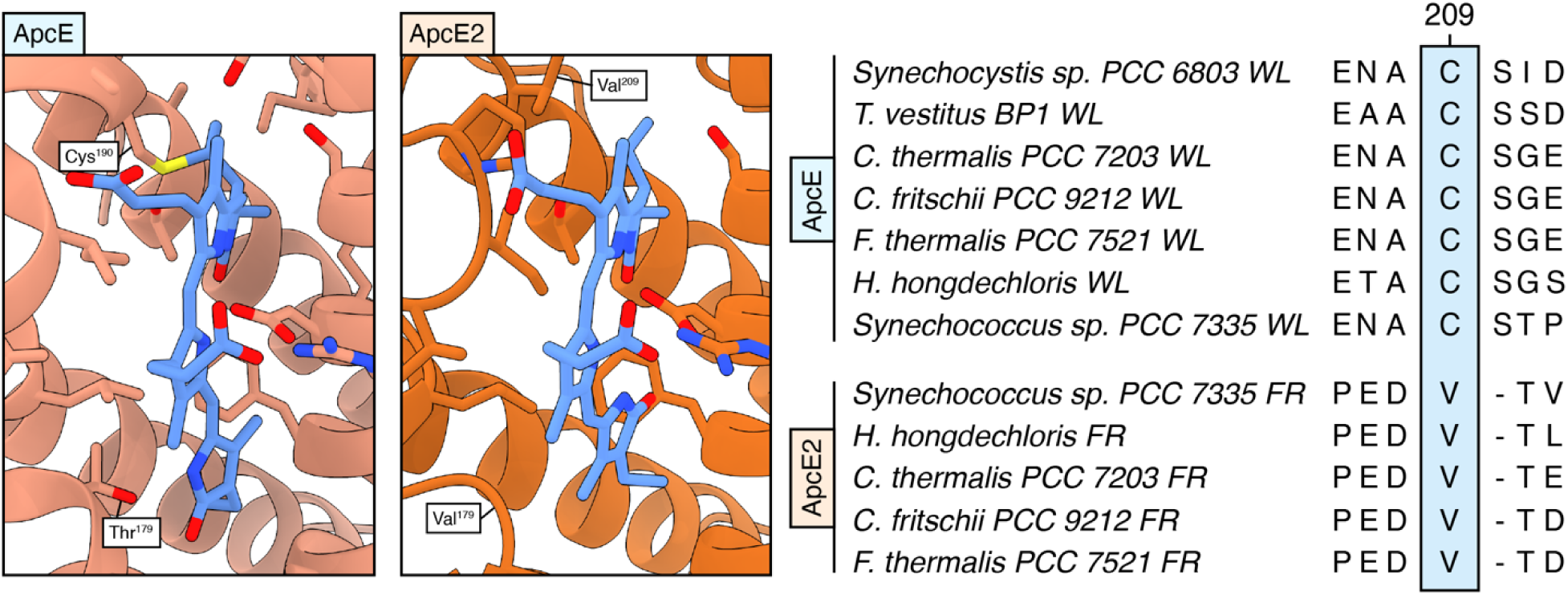
Protein environment of the bilin in the ApcE linker protein. A) Comparison of the chemical environment of the bilin pigment in the WL-ApcE protein (7SC7) against the FR-ApcE2 protein (right panel). The bilin pigment is represented in blue and the protein backbone in orange. B) Sequence alignment between selected ApcE and ApcE2 sequences. The light blue box highlights the discussed conserved loss of Cys^209^ between WL and FR paralogues.

**Fig. 5.**
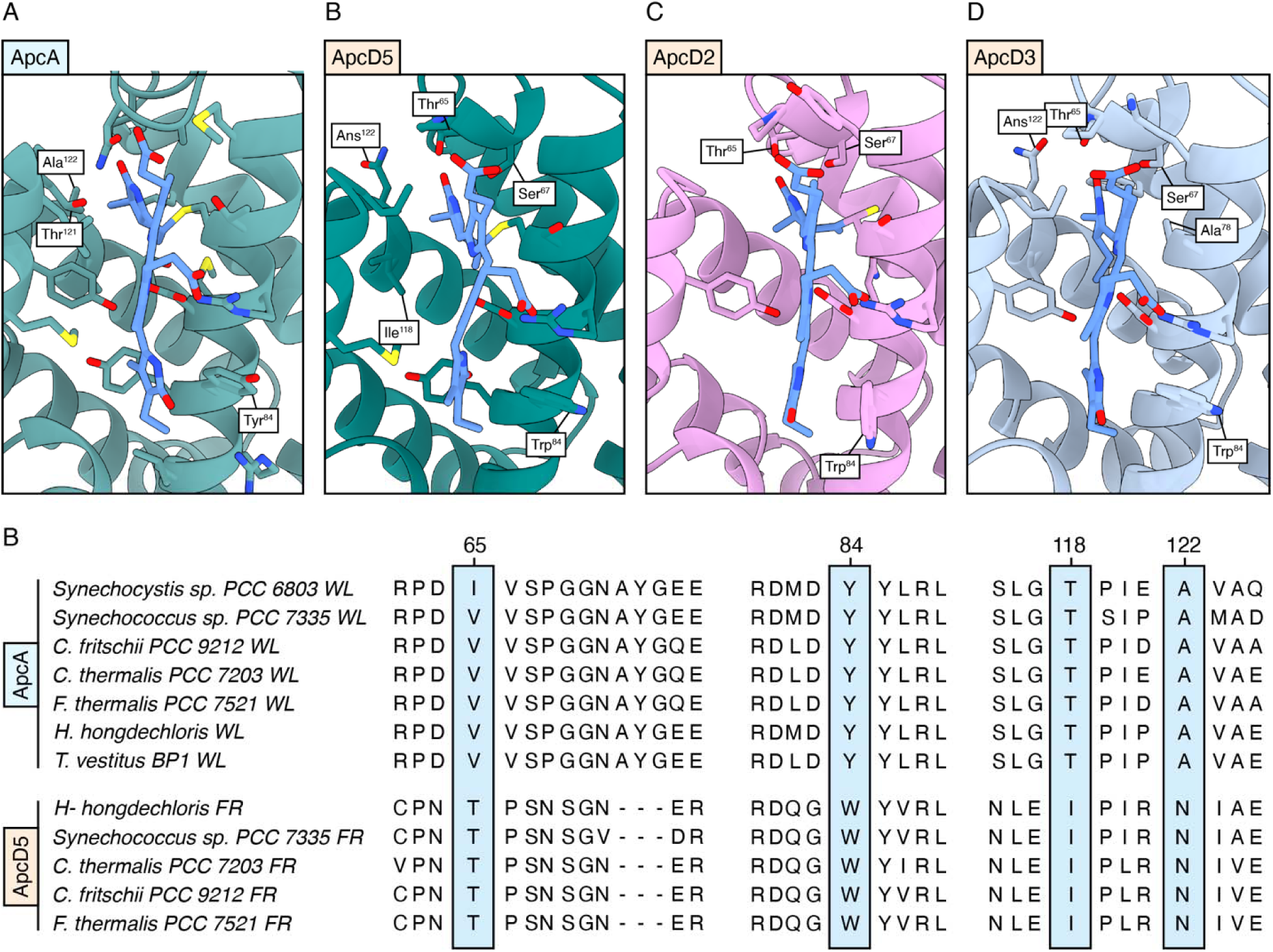
Changes in the bilin sites of ApcD2, ApcD3 and ApcD5 compared with the WL paralogues. Comparison of the chemical environment of the bilin pigment in the WL-ApcA protein (A) against the ApcD5 (B), ApcD3 (C), and ApcD2 (D) proteins. The bilin pigments are always represented in blue. E) Sequence alignment between ApcA and ApcD5, the light blue box highlights conserved changes in the sequence between WL and FR paralogues.

Comparison of far-red protein sequences against the corresponding visible light homologues also revealed amino acid residues that are likely involved in redshifting the absorption spectrum of the complex. The phycocyanobilin in ApcD5, when compared to the one in WL-ApcA, has its rings A and D slightly shifted. This bilin binding is located next to a protein loop which is very different from the WL paralogue (Fig. 5). This loop shifts the position of ring A and provides extra hydrogen bonds via the FR-specific and conserved ApcD5 Thr^65^ and its hydrogen bonding partner ApcD5 Asn^122^ (Fig. 5B). The FR-specific ApcD5 Ile^118^ (Thr in ApcA) also seems to contribute to the shift of ring A by steric hinderance with its longer side chain. The shift of ring D between WL-ApcA and FR-ApcD5 could be attributed to the change from ApcD5 Trp^84^ to Tyr^84^ in ApcA (Fig. 5A, B), resulting in a 0 to ∼30° degrees distortion of ring D relative to ring C (Fig. 3B, Fig. 5B). This same Trp/Tyr change and the ring distortion effects are also seen in ApcD2, another FR-specific α-subunit (Fig. 5C).

For ApcD3, one of the previously proposed emitters (Ho et al., 2017; Gisriel et al., 2024), a significant shift of Ring D is observed compared to its WL paralogue ApcA. The position of ring D of ApcD3 is also influenced by a conserved Thr^75^ to Cys^75^ change in ApcB2. The bulky sulfur side chain of ApcB2 Cys^75^ shifts ring D of the bilin in ApcD3 making it more coplanar with ring C, thereby red-shifting it. A loop between two helices that form part of the ApcD3 bilin site is also changed, resulting in changes in packing and a slight shift of ring C (Fig. 5).

In the far-red specific membrane linker protein ApcE2, the bilin conformation is different from WL-ApcE. The cysteine linkage is lost, the bilin rings are more coplanar, and ring D is rotated 180° degrees compared to that of ApcE (Fig. 4A). This is related to the presence of a conserved Thr^179^ to Val^179^ change (Fig. 4A), that results in the loss of a hydrogen bond to ring D and the consequent flip. All of these changes contribute to the significant redshift of the pigment, consistent with it being the terminal emitter as proposed (Soulier, Laremore & Bryant, 2020; Gisriel et al., 2024).

No major structural changes are observed in the bilin chemical environment of the FR adapted β-subunit ApcB2. Spectroscopically, the absorption spectrum of this subunit is very similar to that of the WL-paralogue ApcB (Ho et al., 2017).

As pointed out in the structural overview of the complex, the distal ApcC subunit, which in WL-APC is present between the third and fourth ring of the complex (Domínguez-Martín et al., 2022), is absent in FR-APC (Fig. 6). In ApcE2 sequences, the loop interacting with the distal ApcC presents poor sequence conservation and some of the residues with bulkier side chains create steric hindrance that would prevent ApcC from inserting in the distal site when compared with the white light paralogue ApcE (Supplementary fig. 2).

**Fig. 6.**
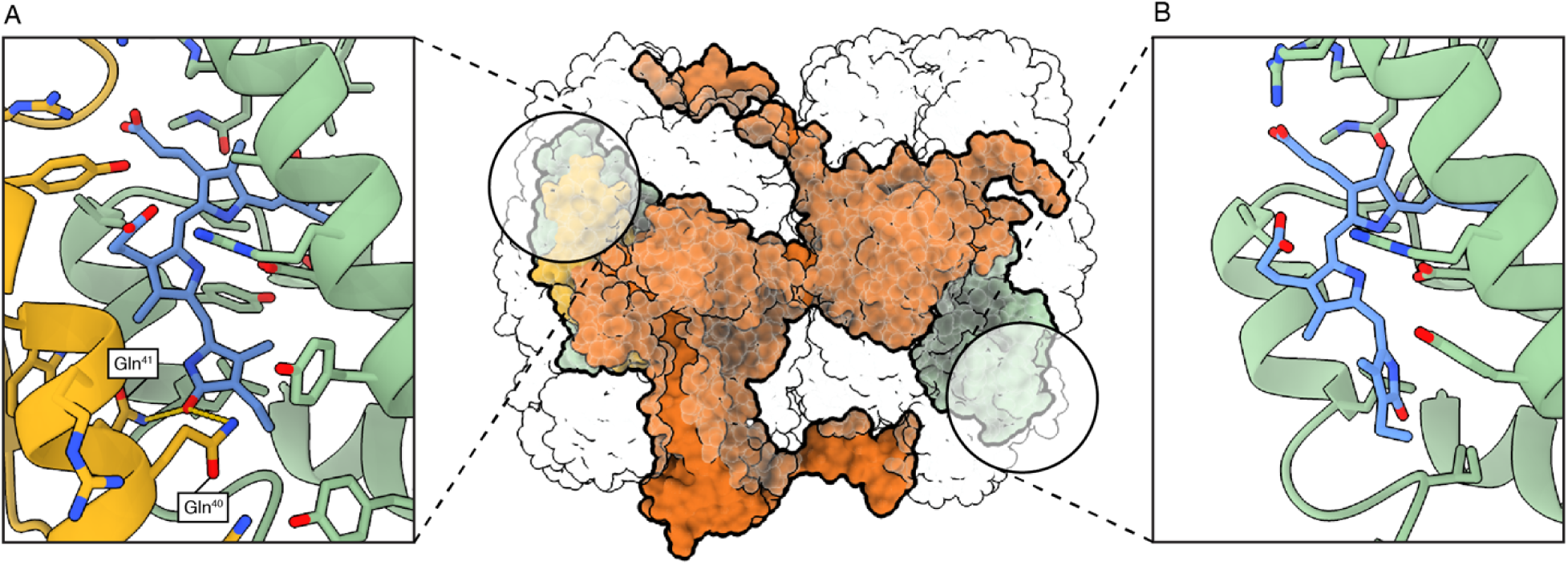
Comparison between the chemical environments of different ApcB2 subunits in the FR-APC complex. In the center, one cylinder with ApcE2 is represented in orange, ApcC is represented in yellow and two ApcB2 subunits in green. A) Proximal ApcB2 bilin binding site in the presence of ApcC. ApcB2 is represented in green, ApcC is represented in yellow, and phycocyanobilin is represented in blue. B) Distal ApcB2 bilin binding site in the absence of ApcC. ApcB2 is represented in green while phycocyanobilin is represented in blue.

This observation suggests that the absence of ApcC is native rather than a purification artifact. By comparing the two relevant ApcB2 subunits, one in the first ring which interacts with ApcC both in WL-APC and FR-APC (Fig. 6A) and one in the fourth ring that interacts in ApcC in WL-APC but not in FR-APC (Fig. 6B), it’s clear that ApcC can radically influence the conformation of bilin pigments. The presence of two hydrogen bonding residues, ApcC Gln^40^ and ApcC Gln^41^, ensure that the orientation of ring A is fixed and rotated by almost 180° compared to the ApcB2 bilin in the absence of ApcC (Fig. 6). Moreover, the bilin pigment is slightly stretched by the presence of these hydrogen bonds, which is likely to affect the absorbance spectrum. The absence of the distal ApcC may tune the pigments in the distal fourth (αβ)_3_ trimer blue-shifting them, creating an energy gradient that facilitates excitation energy transfer to the terminal emitter ApcE2.

The dimer interface of the two cylinders is consistent with that predicted from the previous structural data of the monomeric first three rings of the complex (Gisriel et al., 2024), with ApcD2 being one of the main contributors by interacting extensively with an ApcB2 and an ApcD5 subunit of the third ring, and an ApcD5 subunit of the fourth ring in the neighboring cylinder. Nonetheless, contributions to the structural integrity of the bicylindrical complex are not limited to this subunit. The WL-shared subunit ApcF presents an interaction with ApcF in the opposing cylinder. When the two cylinders are brought in proximity by the dimerization, the two ApcF subunits are in position for long polar sidechains to interact and provide stability to the complex. The C-terminal tail of the membrane linker protein, ApcE2, interacts extensively with an ApcD5 subunit in the third ring of the opposing cylinder (Fig. 1, fig S3).

## Discussion

This work provides the intact structure of the FR-APC bicylindrical antenna complex. The bilin pigment in ApcE2 is non-covalently bound and almost completely planar, pushing the physical properties of phycocyanobilin towards the maximum red-shift attainable by this pigment (Staheli et al., 2021; Miao et al., 2016). Conversely the bilin in ApcD3 presents a less planar structure, suggesting a relative blue-shift compared with ApcE2. The analysis performed on the planarity of the bilin chromophores confirms that, as expected, ApcB2 contains the least planar pigments, together with the fewest amino acid differences, which likely explains why its absorption profile is similar to its WL paralogue.

The structure of the bicylindrical FR-APC prompts questions on the importance of antenna size in far-red light photoacclimation. The FR-APC bicylindrical complex provides an extensive enhancement of the absorption cross-section in the far-red for FR-PSII, providing 24 far-red absorbing bilin pigments compared to the 10 long-wavelength pigments found in a FR-PSII dimer. Nonetheless, as previously pointed out, this is a much lower number compared to the amount of bilin pigments provided, for example, by hemidiscoidal phycobilisome structures that contain hundreds of chromophores (Bryant & Gisriel, 2024). This evolutionary adaptation indicates that space optimization within and between the membranes is preferred over antenna size in far-red adapted thylakoids. This suggestion is supported by the increased thylakoid (and presumably photosystem) density in far-red acclimated cells (MacGregor-Chatwin et al., 2023; Li et al., 2016).

Moreover, the differences in absorption spectra between far-red α (ApcD5) and β (ApcB2) subunits imply that the spectral range covered by the FR-APC evolved to be broader than the WL-PBS complex, even though WL-PBS are bigger and structurally more complex. The steeper absorption gradient of the bilins in FR-APC relative to WL-APC may result in a more efficient excitation transfer from FR-APC to FR-PSII, consistent with the enhanced trapping rate of FR-PSII when the FR-APC is connected (Mascoli et al., 2022).

The alterations during far-red light photoacclimation are not limited to the protein and pigment compositions of antenna proteins and photosystems. The ultrastructure changes in thylakoid organization during FaRLiP reflect the need in these conditions to maximize photosynthetic productivity with the decreased number of photons available. To gain insight on the distances between bilin pigments in the bicylindrical FR-APC and chlorophyll *f* molecules present in FR-PSII, the structural models of FR-APC and FR-PSII (Gisriel et al., 2023) were fitted to the *in situ* CryoEM single particle analysis structure of the PBS-PSII supercomplex from the cyanobacterium *Arthrospira platensis* (Zhang et al., 2024) (Fig. 7A).

**Fig. 7.**
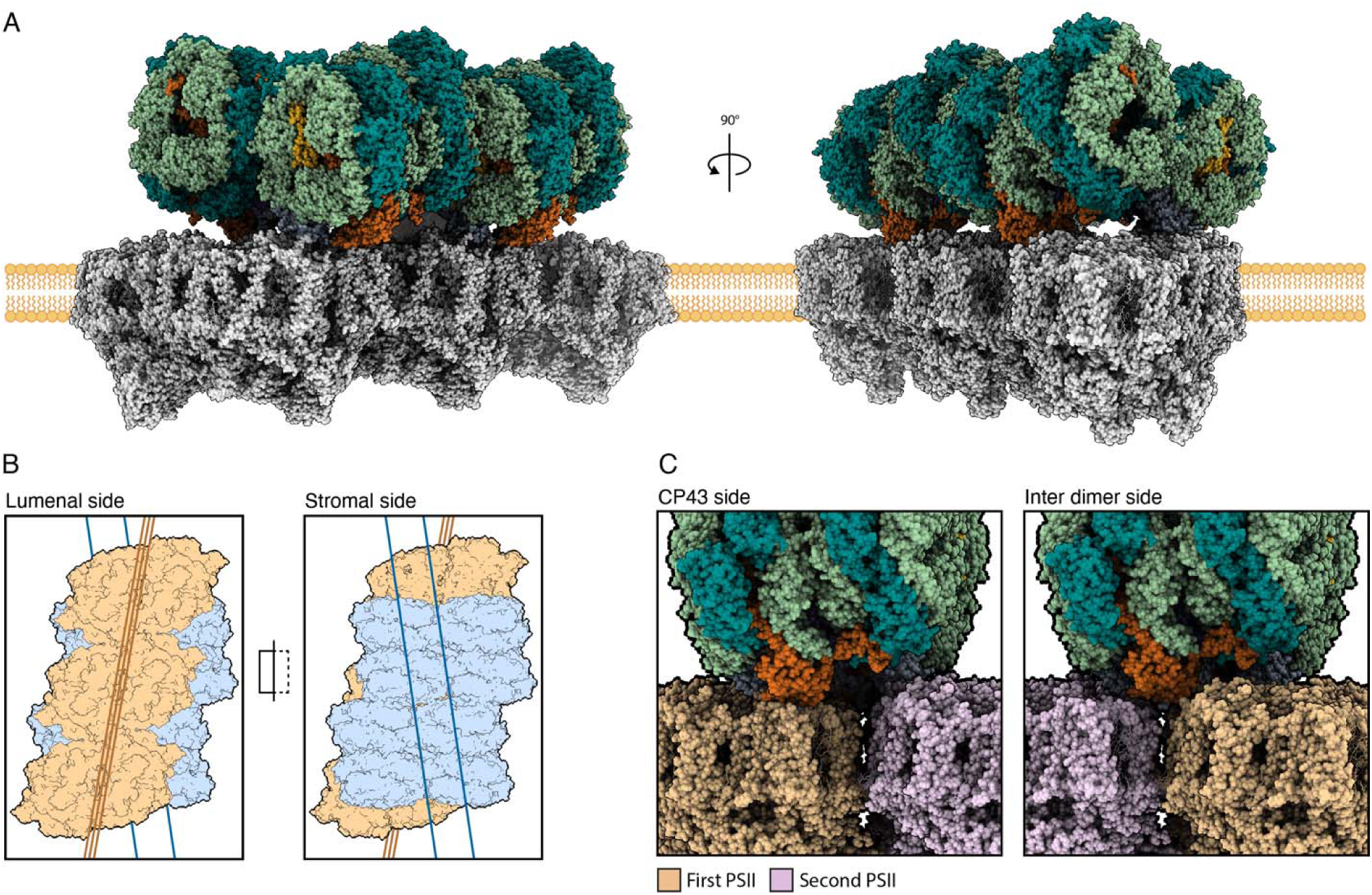
Possible mesoscale arrangement of FR-PSII and bicylindrical FR-APC in thylakoid membranes. A) Representation of an array of three FR-PSII dimers connected to two bicylindrical FR-APC. The representation is modeled based on the arrangement of the PBS-PSII supercomplex from the cyanobacteria *A. platensis* (PDB 8WQL), FR-PSII from *Synechococcus sp.* PCC 7335 (PDB 8EQM) with the solved intact FR-APC. B) Top and bottom view of the FR-APC + FR-PSII array complex with their respective C2 symmetry axes. FR-PSII is represented in orange, while FR-APC is in blue. C) Close-up of the connection of FR-APC to FR-PSII on both sides of the FR-PSII array. In the left panel the interaction with CP43 can be seen and in the right panel ApcE2 is instead between two FR-PSII dimers on the opposite side. FR-APC is colored with the same color scheme as Fig. 1. The two adjacent FR-PSII dimers are represented in orange and pink.

In the model obtained, the symmetry axes of FR-PSII and bicylindrical FR-APC are not parallel but at an angle of roughly ∼20°, suggesting that the two cylinders of the FR-APC are likely to have two independent docking sites on FR-PSII dimers (Fig. 7B) (Zhang et al., 2024). This is confirmed when comparing the two opposite connection sites of the FR-APC cylinders to the array of FR-PSII complexes. On one side, ApcE2 is positioned directly above CP43, while on the other, ApcE2 is wedged between two adjacent FR-PSII dimers. The resolution of the *A. platensis* PBS-PSII supercomplex does not allow the identification of specific residues that stabilize these interactions, and different species may have different interaction interfaces. For this reason, the model serves as a rough indication of distances rather than an actual representation of the array structures in far-red photoacclimated organisms.

Nevertheless, in this model, the closest assigned Chl *f* pigment in FR-PSII to the terminal emitter bilin in ApcE2 is C507. Chl *f* C507 is at an edge-to-edge distance of ∼31 Å from the ApcE2 bilin directly above CP43, and is ∼36 Å away from the ApcE2 bilin between the two dimers, consistent with previous structural and functional studies (Gisriel et al., 2024; Mascoli et al., 2022). Moreover, the bilin pigment contained in ApcD3, previously proposed as a potential terminal emitter in FR-APC, is located further away from any assigned Chl *f* in FR-PSII when compared with the bilin in ApcE2.

While the docked model is not detailed enough to provide precise coordinates, it indicates that phycobilisome complexes, both in WL and FR organisms, are only capable of connecting to PSII when in an ordered array. In such an arrangement, it is possible that excitation energy transfer networks involve transfer between different bicylindrical FR-APC complexes, so that excitation energy can be shared over the entire array of photosystems. Moreover, it is possible that the formation of the array themselves is induced by the presence of ApcE2, stabilizing the side-by-side interaction of adjacent FR-PSII dimers.

From an experimental point of view, this behaviour could explain the inability to resolve structures of APC-PSII complexes via CryoEM single particle analysis, since membrane solubilization treatments would likely destroy the supercomplexes and consequently cause the disconnection of the allophycocyanin cores from the photosystems. This highlights the importance of following new strategies to characterize these complexes, with cryo-electron tomography being an obvious approach (Zhang et al., 2024).

The current work highlights the importance of understanding Chl *f*-based photosynthesis at a scale that goes beyond the individual complexes and their interactions, extending to the organization of the thylakoid membranes and *in situ* photochemistry. Investigating excitation energy transfer networks in the mesoscale will lead to a better understanding of the relationship between antenna size, photosynthetic efficiency, and photodamage when comparing different evolutionary adaptations to far-red light and other ecological niches.

This study illustrates the evolutionary trade-offs that allow efficient management of excitation energy transfer by far-red light adapted antenna. The insights obtained should help with the assessment of the feasibility, requirements, and limitations in the development of applications of far-red photosynthesis.

## Methods

### Culture growth

*Chroococcidiopsis thermalis* PCC 7203 was grown in liquid BG11 medium (Rippka et al., 1979) at 30 °C under far-red light LEDs (750 nm, Epitex; L750-01AU) at an intensity of ∼30 μmol photons m^−2^ s^−1^.

### Isolation of far-red adapted APC+PSII complexes

All purification steps were conducted in the dark and at 4 °C. Cells were harvested and washed in 20mM MES, pH 6.5 buffer containing protease inhibitors (SIGMAFAST protease inhibitor tablets). Cells were broken with two passages in a continuous flow cell disruptor (Constant Systems) at a pressure of 39 kPsi and then centrifuged at 1000x *g* to remove any unbroken material. The supernatant was subsequently centrifuged for 20 minutes at 40,000 rpm in a Ti45 rotor at 4° C to pellet the membranes. Membranes were resuspended in low salt buffer A (20 mM MES, pH 6.5, 5 mM MgCl_2_, 5 mM CaCl_2_, 1.2 M betaine, 0.03% β-DM), adjusted to a chlorophyll concentration of 0.4 mg/ml and solubilized at 4° C for 1 hour by the addition of β-DM to a final concentration of 0.4% (w/v). Solubilized membranes were centrifuged for 30 minutes at 42,000 rpm at 4 °C in Ti45 rotor to remove unsolubilized material. Isolation of protein complexes was carried out with a modified protocol based on Kern et al. (Kern et al., 2005). The solubilized supernatant was loaded onto a DEAE Toyopearl 650S (ID: 50 mm Length: 450 mm), washed with 1 column volume of buffer A and eluted with a 0%-100% linear gradient of buffer B (20 mM MES, pH 6.5, 5 mM MgCl_2_, 5 mM CaCl_2_, 0.5 M NaCl, 1.2 M betaine, 0.03% β-DM) over 5 column volumes. Elution fractions containing protein complexes were concentrated (100 kDa or 50 kDa Filters, Amicon), washed with buffer A and stored at 4 °C or -80 °C depending on the use.

### Grid preparation

Gold quantifoil (2 μm hole size and 1 μm hole spacing) with a 300-copper mesh grids (Quantifoil Micro Tools GmbH) were glow-discharged for 30 s at 25 mA. APC+PSII sample (∼4.2 mg/ml of protein) was applied for 3 seconds at a temperature of 4 °C and 100% humidity in the presence of only dim green light, and the grids were blotted and plunge-frozen in liquid ethane with a Vitrobot mark IV (Thermo Fisher Scientific).

### Data acquisition

The particles were imaged using a Krios 3 operated at 300 kV. Images were recorded on a Falcon 4i with a pixel size of 0.723 Å and an exposure of 40 electrons per Å^2^ for a total of 40 frames. Images were collected in super resolution mode with a SelectrisX energy filter with a slit width of 20 eV. The targeted defocus range was varied from –0.6 to –2.4 µm using the EPU software (Thermo Fisher).

### Single particle analysis

A total of 9397 movies were collected. Frames were aligned, dose weighted, and the contrast transfer function (CTF) was estimated in CryoSPARC v4.0.2 (Punjani et al., 2017).

Micrographs were curated by removing ones with CTF fits below 10 Å. The subset obtained contained 9224 micrographs (98%) and blob picker was used to pick particles in a subset of micrographs across the defocus range. Particles were 2D classified and used to train Topaz (Bepler et al., 2019) to pick particles across the entire dataset, yielding 423,203 picks. After multiple rounds of 2D classification and *ab-initio* refinement, duplicated particles were removed and a subset of 36,300 particles were used for homogeneous refinement to obtain an initial map >5Å and to confirm C2 symmetry. After multiple rounds of per particle CTF refinement and local motion correction (Rubinstein & Brubaker, 2015), a set of 26,300 particles were used to perform non-uniform refinement (Punjani, Zhang & Fleet, 2020) imposing C2 symmetry, producing a map at a global resolution of 2.61 Å based on GC-Fourier Shell Correlation at a cut-off of 0.143.

### Model building

The FR-APC incomplete rod from 8UHE (Gisriel et al., 2024) was fitted to the ESP map using the Phenix software suite (Adams et al., 2010), and then the third ring was copied and fitted again with Phenix into the density of the fourth ring. The complete model was then mutated to the correct sequence with Chainsaw (CCP4) and refined in Coot (Emsley et al., 2010). The amino acid for each subunit’s density was evaluated individually and the model was then refined in real space in Phenix (table S2).

### Absorbance measurements

Absorption spectra of isolated complexes were recorded at room temperature with a Cary 60 spectrophotometer (Agilent technologies) in buffer A.

### 77K fluorescence spectroscopy

Low temperature fluorescence measurements were performed on isolated FR-PSII + FR-APC complexes at an OD_710_ of 0.1 with a FluoroMax 4 spectrofluorometer (HORIBA Ltd., Japan). 77K fluorescence emission was measured from 650-850 nm with excitation and emission slits of 2 nm and an integration time of 0.5 s using an excitation range from 400-600 nm.

## Acknowledgments

We thank Kenta Renard for the numerous discussions. We thank the centre for structural biology at Imperial College for the training provided, for help with the early stages of sample screening and with data collection. We thank Diamond for the access and support of the cryo-EM facilities at the UK national electron Bio-Imaging Centre (eBIC), proposal BI25127.

## Funding

Funding information, including grant numbers, complete funding agency names, and recipient’s initials:

Marie Skłodowska-Curie grant agreement, No. 955520 – GC, AWR, AF

Biotechnology and Biological Sciences Research Council, BB/R001383/1 – AWR, AF, GAD

Biotechnology and Biological Sciences Research Council, BB/V002015/1 – AWR, AF, JWM

Biotechnology and Biological Sciences Research Council, BB/R00921X – AWR, AF, GAD

Leverhulme Trust Grant RPG-2022-203 – AWR, GAD, AF

The Royal Society (Royal Society Research Professorship 2024) - AWR

## Author contributions

GC, HFL initiated the study.

GAD grew and harvested *C. thermalis* PCC 7203

GC, HFL, GAD, AF, and AWR conceived of the main experiments, collated results and interpretations, and wrote the article with input, edits, and approval from all authors.

GAD developed and performed the isolation of the complexes.

GC and HFL prepared the grids.

GC and HFL collected the micrographs.

GC and HFL processed the data to obtain the map.

GC, HFL and JWM built the atomic model.

HFL collected the sequences and performed the phylogenetic analysis.

GC coded the bilin planarity quantification method.

GC prepared the figures with input, edits, and approval from all other authors

## Data and materials availability

The atomic coordinates have been deposited in the PDB with accession code 9I1R. The script used to calculate and visualize bilin angles is available upon request to the authors.

## Supplementary Materials

Fig. S1 to S3

Tables S1 and S2

## Bibliography

1. Adams, P.D., Afonine, P. V, Bunkóczi, G., Chen, V.B., Davis, I.W., Echols, N., Headd, J.J., Hung, L.-W., Kapral, G.J., Grosse-Kunstleve, R.W., McCoy, A.J., Moriarty, N.W., Oeffner, R., Read, R.J., Richardson, D.C., Richardson, J.S., Terwilliger, T.C. & Zwart, P.H. (2010) PHENIX: a comprehensive Python-based system for macromolecular structure solution. Acta Crystallographica Section D. 66 (2), 213–221. doi:10.1107/S0907444909052925.

2. Antonaru, L.A., Cardona, T., Larkum, A.W.D. & Nürnberg, D.J. (2020) Global distribution of a chlorophyll f cyanobacterial marker. The ISME Journal. 14 (9), 2275–2287. doi:10.1038/s41396-020-0670-y.

3. Behrendt, L., Trampe, E.L., Nord, N.B., Nguyen, J., Kühl, M., Lonco, D., Nyarko, A., Dhinojwala, A., Hershey, O.S. & Barton, H. (2020) Life in the dark: far-red absorbing cyanobacteria extend photic zones deep into terrestrial caves. Environmental Microbiology. 22 (3), 952–963. doi:10.1111/1462-2920.14774.

4. Bepler, T., Morin, A., Rapp, M., Brasch, J., Shapiro, L., Noble, A.J. & Berger, B. (2019) Positive-unlabeled convolutional neural networks for particle picking in cryo-electron micrographs. Nature Methods. 16 (11), 1153–1160. doi:10.1038/s41592-019-0575-8.

5. Brejc, K., Ficner, R., Huber, R. & Steinbacher, S. (1995) Isolation, crystallization, crystal structure analysis and refinement of allophycocyanin from the cyanobacterium Spirulina platensis at 2.3 Å resolution. Journal of molecular biology. 249 (2), 424–440.

6. Bryant, D.A. & Gisriel, C.J. (2024) The structural basis for light harvesting in organisms producing phycobiliproteins. The Plant Cell. koae126.

7. Bryant, D.A., Glazer, A.N. & Eiserling, F.A. (1976) Characterization and structural properties of the major biliproteins of Anabaena sp. Archives of Microbiology. 110, 61–75.

8. Bryant, D.A., Shen, G., Turner, G.M., Soulier, N., Laremore, T.N. & Ho, M.-Y. (2020) Far-red light allophycocyanin subunits play a role in chlorophyll d accumulation in far-red light. Photosynthesis research. 143, 81–95.

9. Chen, M., Schliep, M., Willows, R.D., Cai, Z.-L., Neilan, B.A. & Scheer, H. (2010) A red-shifted chlorophyll. Science. 329 (5997), 1318–1319.

10. Domínguez-Martín, M.A., Sauer, P. V., Kirst, H., Sutter, M., Bína, D., Greber, B.J., Nogales, E., Polívka, T. & Kerfeld, C.A. (2022) Structures of a phycobilisome in light-harvesting and photoprotected states. Nature 2022 609:7928. 609 (7928), 835–845. doi:10.1038/s41586-022-05156-4.

11. Emsley, P., Lohkamp, B., Scott, W.G. & Cowtan, K. (2010) Features and development of Coot. Acta Crystallographica Section D: Biological Crystallography. 66 (4), 486–501. doi:10.1107/S0907444910007493.

12. Gan, F. & Bryant, D.A. (2015) Adaptive and acclimative responses of cyanobacteria to far red light. Environmental microbiology. 17 (10), 3450–3465.

13. Gan, F., Zhang, S., Rockwell, N.C., Martin, S.S., Lagarias, J.C. & Bryant, D.A. (2014) Extensive remodeling of a cyanobacterial photosynthetic apparatus in far-red light. Science. 345 (6202), 1312–1317. doi:10.1126/science.1256963.

14. Gisriel, C.J., Shen, G., Brudvig, G.W. & Bryant, D.A. (2024) Structure of the antenna complex expressed during far-red light photoacclimation in Synechococcus sp. PCC 7335. Journal of Biological Chemistry. 300 (2), 105590. doi: 10.1016/j.jbc.2023.105590.

15. Gisriel, C.J., Shen, G., Flesher, D.A., Kurashov, V., Golbeck, J.H., Brudvig, G.W., Amin, M. & Bryant, D.A. (2023) Structure of a dimeric photosystem II complex from a cyanobacterium acclimated to far-red light. Journal of Biological Chemistry. 299 (1). doi:10.1016/j.jbc.2022.102815.

16. Ho, M.-Y., Gan, F., Shen, G. & Bryant, D.A. (2017) Far-red light photoacclimation (FaRLiP) in Synechococcus sp. PCC 7335. II. Characterization of phycobiliproteins produced during acclimation to far-red light. Photosynthesis research. 131, 187–202.

17. Kern, J., Loll, B., Lüneberg, C., DiFiore, D., Biesiadka, J., Irrgang, K.D. & Zouni, A. (2005) Purification, characterisation and crystallisation of photosystem II from Thermosynechococcus elongatus cultivated in a new type of photobioreactor. Biochimica et Biophysica Acta (BBA) - Bioenergetics. 1706 (1–2), 147–157. doi:10.1016/J.BBABIO.2004.10.007.

18. Li, Y., Lin, Y., Garvey, C.J., Birch, D., Corkery, R.W., Loughlin, P.C., Scheer, H., Willows, R.D. & Chen, M. (2016) Characterization of red-shifted phycobilisomes isolated from the chlorophyll f-containing cyanobacterium Halomicronema hongdechloris. Biochimica et Biophysica Acta (BBA) - Bioenergetics. 1857 (1), 107–114. doi:10.1016/J.BBABIO.2015.10.009.

19. MacGregor-Chatwin, C., Nürnberg, D.J., Jackson, P.J., Vasilev, C., Hitchcock, A., Ho, M.-Y., Shen, G., Gisriel, C.J., Wood, W.H.J., Mahbub, M., Selinger, V.M., Johnson, M.P., Dickman, M.J., Rutherford, A.W., Bryant, D.A. & Hunter, C.N. (2023) Changes in supramolecular organization of cyanobacterial thylakoid membrane complexes in response to far-red light photoacclimation. Science Advances. 8 (6), eabj4437. doi:10.1126/sciadv.abj4437.

20. Mascoli, V., Bhatti, A.F., Bersanini, L., van Amerongen, H. & Croce, R. (2022) The antenna of far-red absorbing cyanobacteria increases both absorption and quantum efficiency of Photosystem II. Nature Communications. 13 (1), 3562. doi:10.1038/s41467-022-31099-5.

21. Miao, D., Ding, W.-L., Zhao, B.-Q., Lu, L., Xu, Q.-Z., Scheer, H. & Zhao, K.-H. (2016) Adapting photosynthesis to the near-infrared: non-covalent binding of phycocyanobilin provides an extreme spectral red-shift to phycobilisome core-membrane linker from Synechococcus sp. PCC7335. Biochimica et Biophysica Acta (BBA)-Bioenergetics. 1857 (6), 688–694.

22. Nürnberg, D.J., Morton, J., Santabarbara, S., Telfer, A., Joliot, P., Antonaru, L.A., Ruban, A. V, Cardona, T., Krausz, E., Boussac, A., Fantuzzi, A. & Rutherford, A.W. (2018) Photochemistry beyond the red limit in chlorophyll f–containing photosystems. Science. 360 (6394), 1210–1213. doi:10.1126/science.aar8313.

23. Ohkubo, S. & Miyashita, H. (2017) A niche for cyanobacteria producing chlorophyll f within a microbial mat. The ISME Journal. 11 (10), 2368–2378. doi:10.1038/ismej.2017.98.

24. Punjani, A., Rubinstein, J.L., Fleet, D.J. & Brubaker, M.A. (2017) cryoSPARC: algorithms for rapid unsupervised cryo-EM structure determination. Nature Methods. 14 (3), 290–296. doi:10.1038/nmeth.4169.

25. Punjani, A., Zhang, H. & Fleet, D.J. (2020) Non-uniform refinement: adaptive regularization improves single-particle cryo-EM reconstruction. Nature Methods. 17 (12), 1214–1221. doi:10.1038/s41592-020-00990-8.

26. Rippka, R., Deruelles, J., Waterbury, J.B., Herdman, M. & Stanier, R.Y. (1979) Generic Assignments, Strain Histories and Properties of Pure Cultures of Cyanobacteria. Microbiology. 111 (1), 1–61. doi:10.1099/00221287-111-1-1.

27. Rubinstein, J.L. & Brubaker, M.A. (2015) Alignment of cryo-EM movies of individual particles by optimization of image translations. Journal of Structural Biology. 192 (2), 188–195. doi:10.1016/j.jsb.2015.08.007.

28. Soulier, N. & Bryant, D.A. (2021) The structural basis of far-red light absorbance by allophycocyanins. Photosynthesis research. 147 (1), 11–26.

29. Soulier, N., Laremore, T.N. & Bryant, D.A. (2020) Characterization of cyanobacterial allophycocyanins absorbing far-red light. Photosynthesis research. 145 (3), 189–207.

30. Soulier, N.T. & Bryant, D.A. (2023) Long-wavelength phycobiliproteins. In: Photosynthesis. Elsevier. pp. 9–32.

31. Staheli, C.F., Barney, J., Clark, T.R., Bowles, M., Jeppesen, B., Oblinsky, D.G., Steffensen, M.B. & Dean, J.C. (2021) Spectroscopic and Photophysical Investigation of Model Dipyrroles Common to Bilins: Exploring Natural Design for Steering Torsion to Divergent Functions. Frontiers in Chemistry. 9, 628852.

32. Trampe, E. & Kühl, M. (2016) Chlorophyll f distribution and dynamics in cyanobacterial beachrock biofilms. Journal of phycology. 52 (6), 990–996.

33. Xu, Q.-Z., Tang, Q.-Y., Han, J.-X., Ding, W.-L., Zhao, B.-Q., Zhou, M., Gärtner, W., Scheer, H. & Zhao, K.-H. (2017) Chromophorylation (in Escherichia coli) of allophycocyanin B subunits from far-red light acclimated Chroococcidiopsis thermalis sp. PCC7203. Photochemical & Photobiological Sciences. 16, 1153–1161.

34. Zhang, X., Xiao, Y., You, X., Sun, S. & Sui, S.F. (2024) In situ structural determination of cyanobacterial phycobilisome–PSII supercomplex by STAgSPA strategy. Nature Communications 2024 15:1. 15 (1), 1–11. doi:10.1038/s41467-024-51460-0.

35. Zhao, C., Gan, F., Shen, G. & Bryant, D.A. (2015) RfpA, RfpB, and RfpC are the master control elements of far-red light photoacclimation (FaRLiP). Frontiers in Microbiology. 6, 1303.

36. Zlenko, D. V, Elanskaya, I. V, Lukashev, E.P., Bolychevtseva, Y. V, Suzina, N.E., Pojidaeva, E.S., Kononova, I.A., Loktyushkin, A. V & Stadnichuk, I.N. (2019) Role of the PB-loop in ApcE and phycobilisome core function in cyanobacterium Synechocystis sp. PCC 6803. Biochimica et Biophysica Acta (BBA)-Bioenergetics. 1860 (2), 155–166.

