## Supplementary Materials for "Structure of far-red allophycocyanin: stripped down and tuned up for low energy photosynthesis"

### Title:

### Supplementary materials

|  | FR-APC |
| --- | --- |
| **Data collection** |  |
| Microscope | Krios III |
| Camera | Falcon 4i |
| Magnification | 155.000x |
| keV | 300 |
| e-/Å2 | 40 |
| Defocus range | -0.8 to -2 |
| Å/px | 0.723 |
| Energy filter | Selectris (20 eV) |
| Exposures | 9397 |
| **Data processing** |  |
| Box size | 700 px |
| CTF <10 Å | 0.98 |
| Initial particles | 423204 |
| Final particles | 36300 |
| Symmetry | C2 |
| GS-FSC 0.143 | 2.61 |
| B-factor | 47.9 |

Table S1 - collection and data processing parameters

Table presenting the CryoEM collection parameters and the ones used during Cryosparc processing

| Protein Geometry | Poor rotamers | 1 | 0.01% | Goal: <0.3% |
| --- | --- | --- | --- | --- |
|  | Favored rotamers | 7406 | 99.76% | Goal: >98% |
|  | Ramachandran outliers | 3 | 0.03% | Goal: <0.05% |
|  | Ramachandran favored | 8488 | 98.13% | Goal: >98% |
|  | Rama distribution Z-score | 2.27 ± 0.09 | | Goal: abs(Z score) < 2 |
|  | Cβ deviations >0.25Å | 0 | 0.00% | Goal: 0 |
|  | Bad bonds: | 0 / 72560 | 0.00% | Goal: 0% |
|  | Bad angles: | 3 / 98342 | 0.00% | Goal: <0.1% |
| Peptide Omegas | Cis Prolines: | 4 / 298 | 1.34% | Expected: ≤1 per chain, or ≤5% |
| Low-resolution Criteria | CaBLAM outliers | 77 | 0.9% | Goal: <1.0% |
|  | CA Geometry outliers | 15 | 0.17% | Goal: <0.5% |
| Additional validations | Chiral volume outliers | 0/10838 | |  |

Table S2 – MolProbity statistics

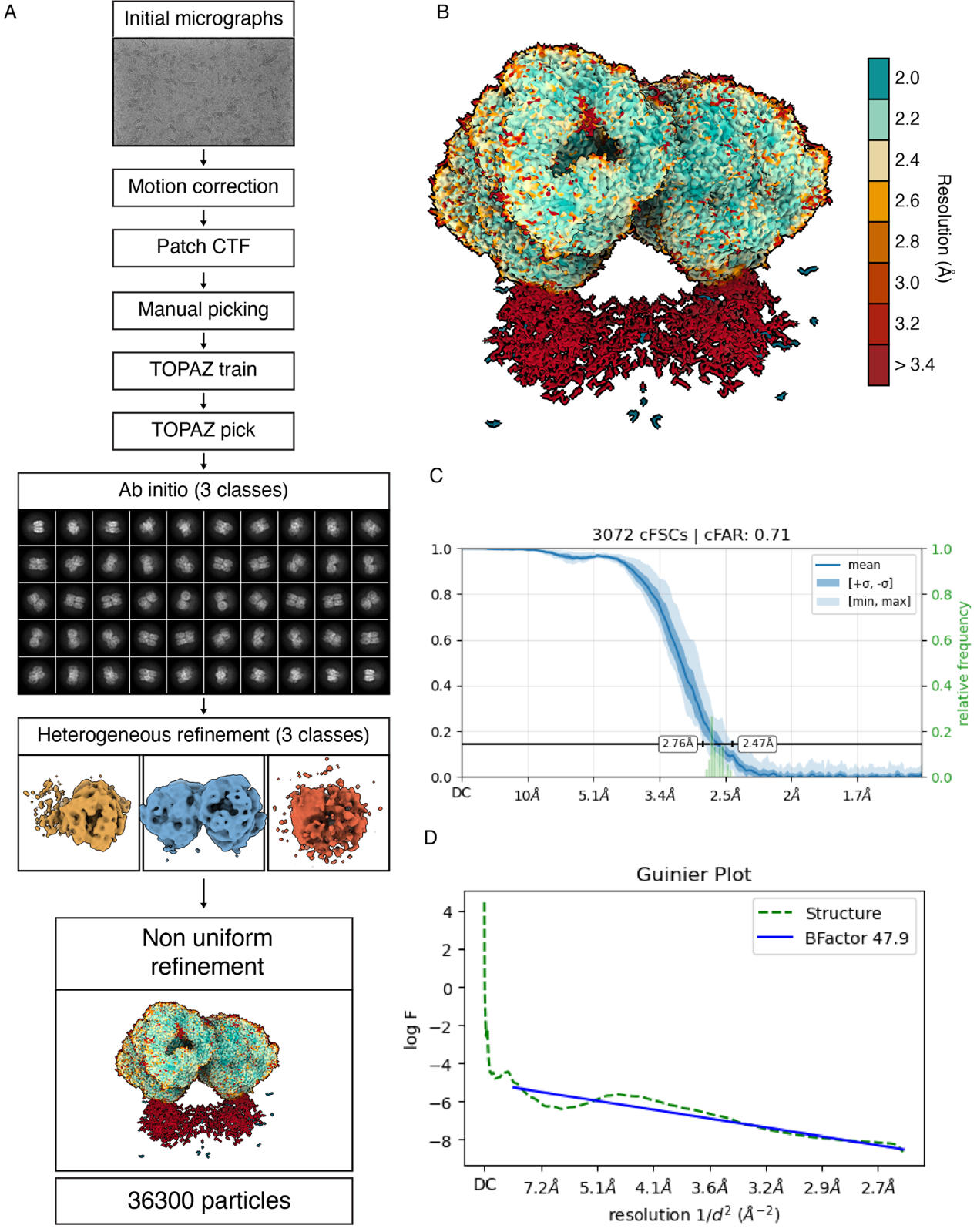

Fig. S1 - CryoSPARC workflow for the SPA structure determination of the FR-APC complex

A) CryoSPARC workflow for the determination of the map of the FR-APC complex. B) local resolution map of the FR-APC complex colored according to the colorbar on the side. C) GS-FSC plot of the resolution of the complex D) Guinier’s plot of the FR-APC complex.

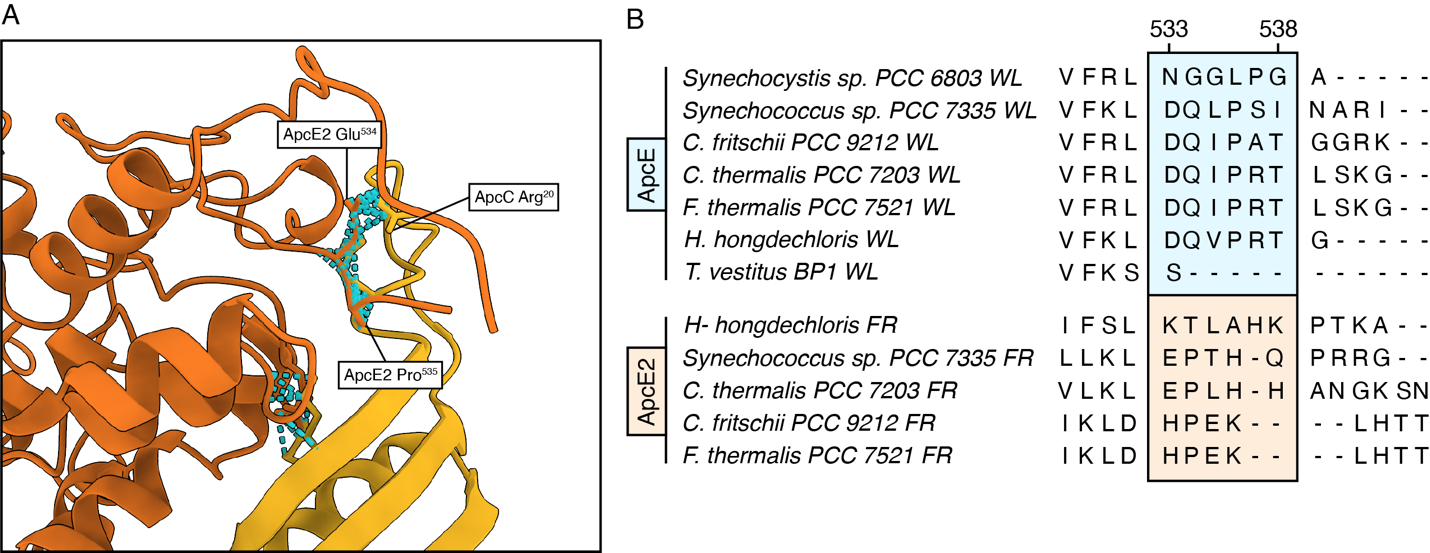

Fig. S2 - Clashes between distal ApcC and ApcE2

A) Structural analysis of the potential interaction between ApcC and ApcE2 if ApcC was in the same position compared to WL-PBS. ApcE2 is represented in orange and ApcC is represented in yellow, in blue are represented the molecular clashes between the two subunits B) phylogenetic analysis of the ApcE and ApcE2 FR and WL. In blue and orange, the areas of non-conservation in the C-terminal are highlighted.

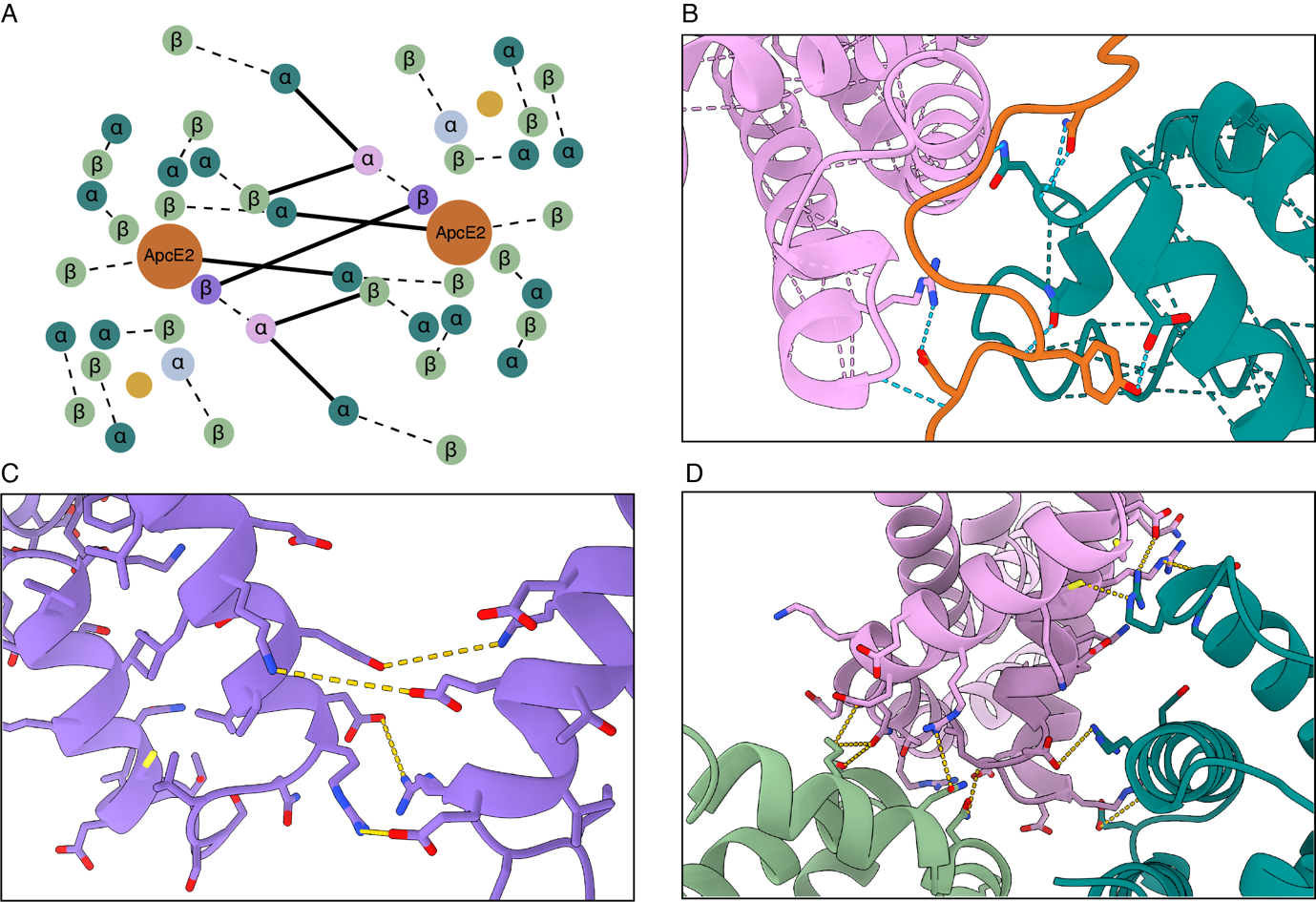

Fig. S3 – Interactions at the dimer interface between the two FR-APC cylinders.

A) Network of the interaction between monomers of the bicylindrical FR-APC. The subunit color key is the same one used throughout the main text. Subunits are represented as circles of dimension proportional to their length in amino acids. Interaction of α and β subunits to form (αβ) monomers is represented as dashed lines, while inter-cylinder interaction is represented by black solid lines. B) Interaction of ApcE2 in orange, with an ApcD5 subunit of the third ring of the opposing cylinder. C) Interaction of the two ApcF from the two different cylinders. D) ApcD2 interactions with an ApcB2 of the third ring and an ApcD5 subunit of the fourth ring of the opposing cylinder.
